## Supporting Information for "Mice with renal-specific alterations of stem cell-associated signaling develop symptoms of chronic kidney disease but surprisingly no tumors"

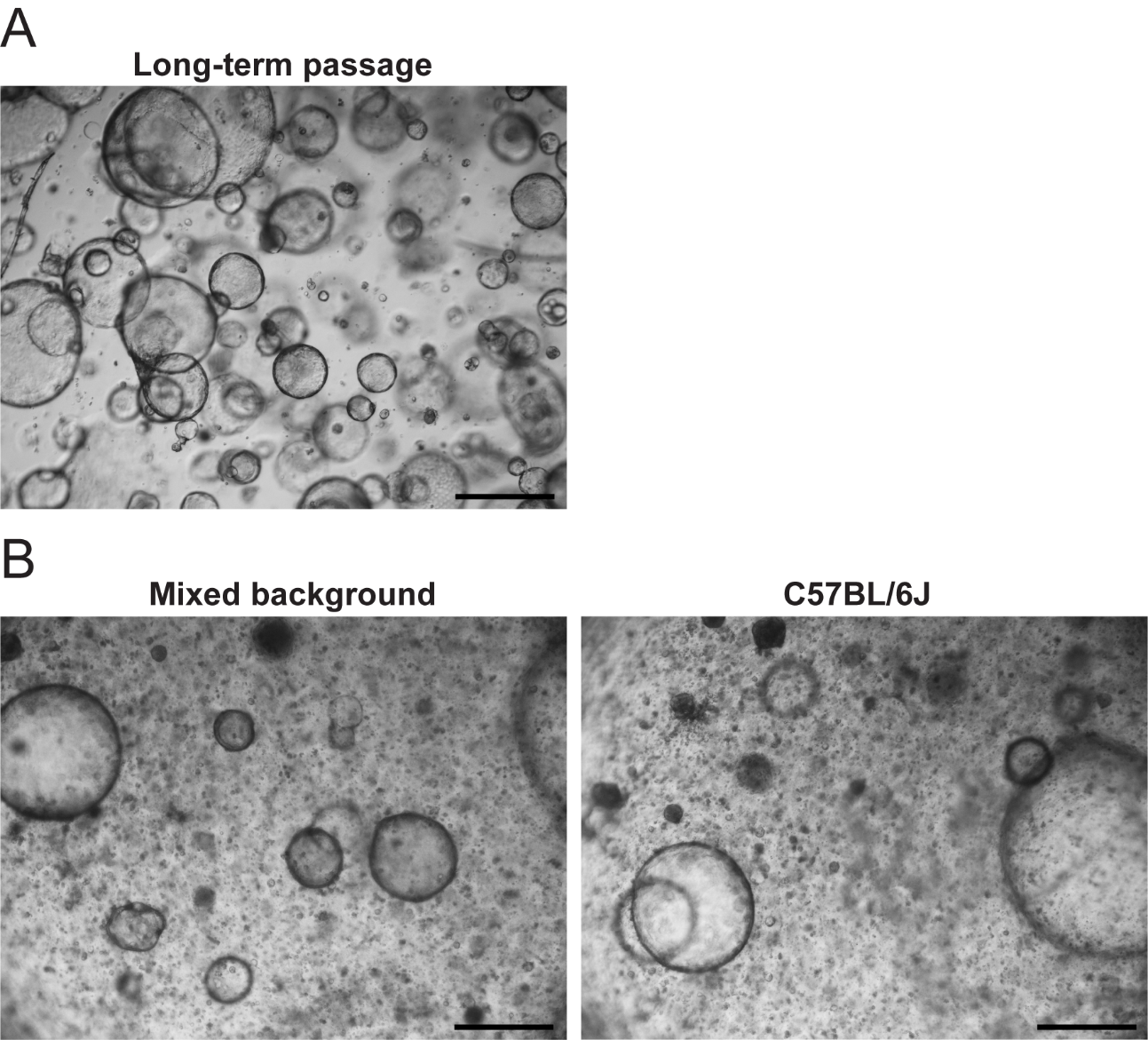

**S1 Figure. Long-term culture of kidney organoids and establishing kidney organoids of different mouse backgrounds.** (A) Brightfield image of an organoid culture, which was serially passaged for 3.5 months. (B) Brightfield image of a freshly seeded organoid culture from a mixed C57BL/6J and FVB/NJ background (left), and from the B57BL/6J background (right) mice. Data information: scale bars, 500 µm. Three independent replicates were examined.

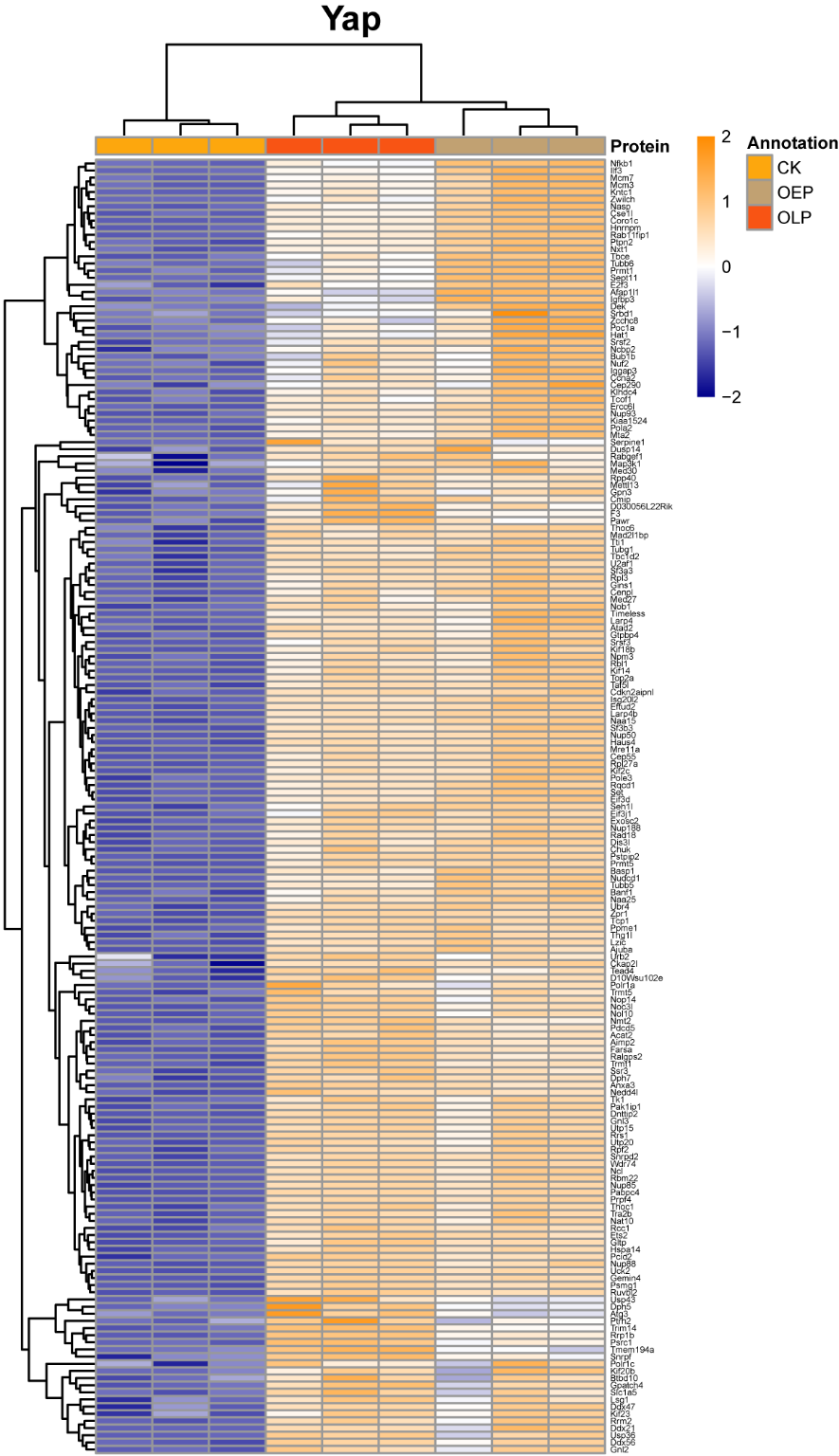

**S2 Figure. Yap signaling is upregulated in kidney organoids.** Proteomic heatmap for direct Yap targets in both early (OEP) and long-term (OLP) passage organoids in comparison to mouse kidney epithelia (control kidney, CK). Data information: the heatmap shows normalized log2 intensity values for three independent replicates of CK, OEP and OLP. A 5% FDR (adjusted P-value < 0.05) cutoff and a log2 fold change cutoff of > 0 were applied for both OEP over CK and OLP over CK. Values were scaled (z-score by row) with breaks from ≤ -2 to ≥ 2.

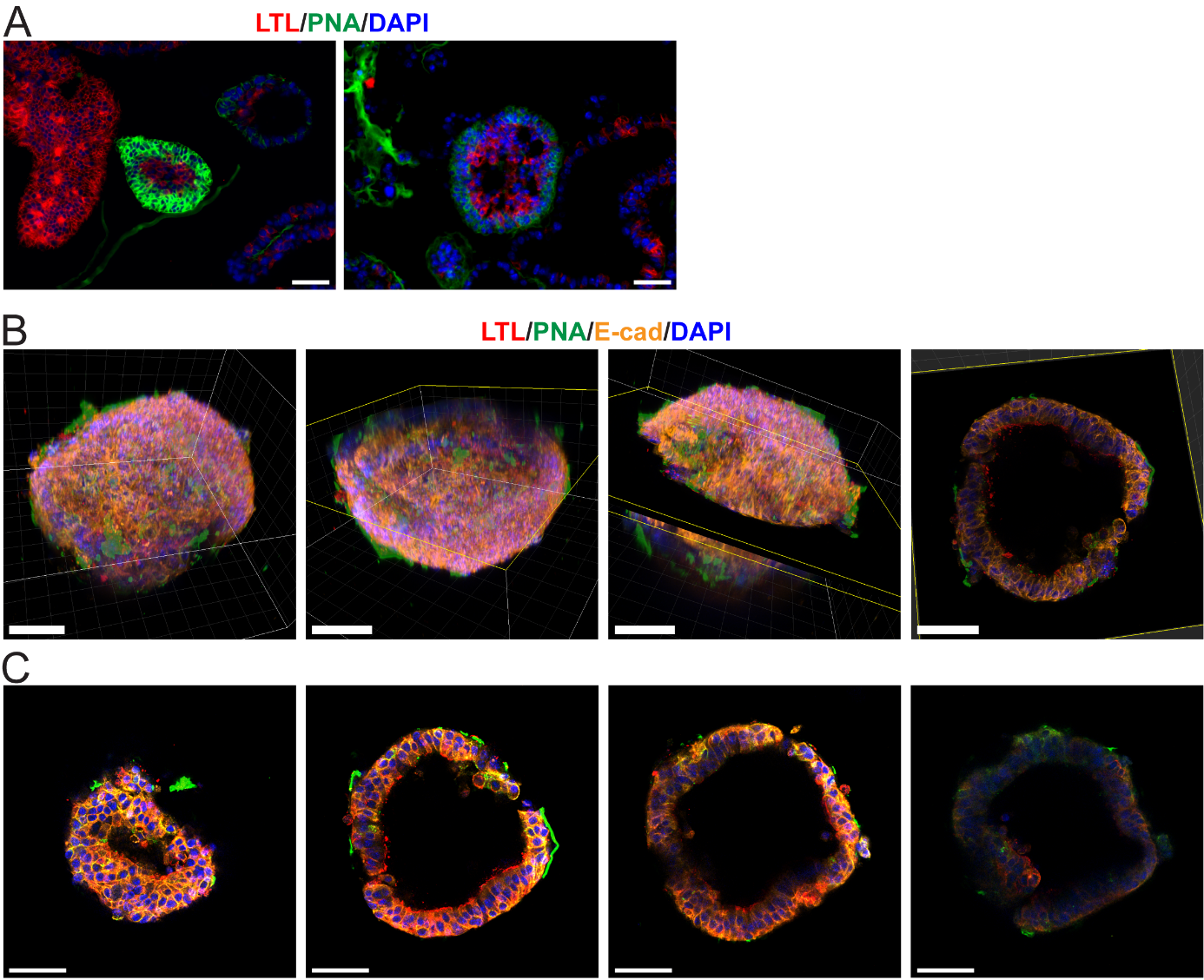

**S3 Figure. Markers of tubular epithelial cells are expressed in 3D reconstructed kidney organoids.** (A) 2D immunofluorescence of a solid (left) and an alveolar (right) organoid positive for both LTL (red) and PNA (green). (B and C) Whole-mount 3D confocal microscopy of a cystic organoid positive for LTL (red), PNA (green) and E-cad (orange). (B) A 3D reconstruction of z-stacks of a cystic organoid. From left to right: a 3D reconstructed organoid seen from the top-side (coordinates given for orientation), top-side view opened by a clipping plane, top-side view with a marked orthogonal slice (8 µm) and extended orthogonal slice (8 µm) only. (C) Selected 2D images of z-stacks at different depths (top, middle, bottom) of a cystic organoid. Data information: scale bars, 50 µm. Nuclei are counterstained with DAPI. Three independent replicates were examined.

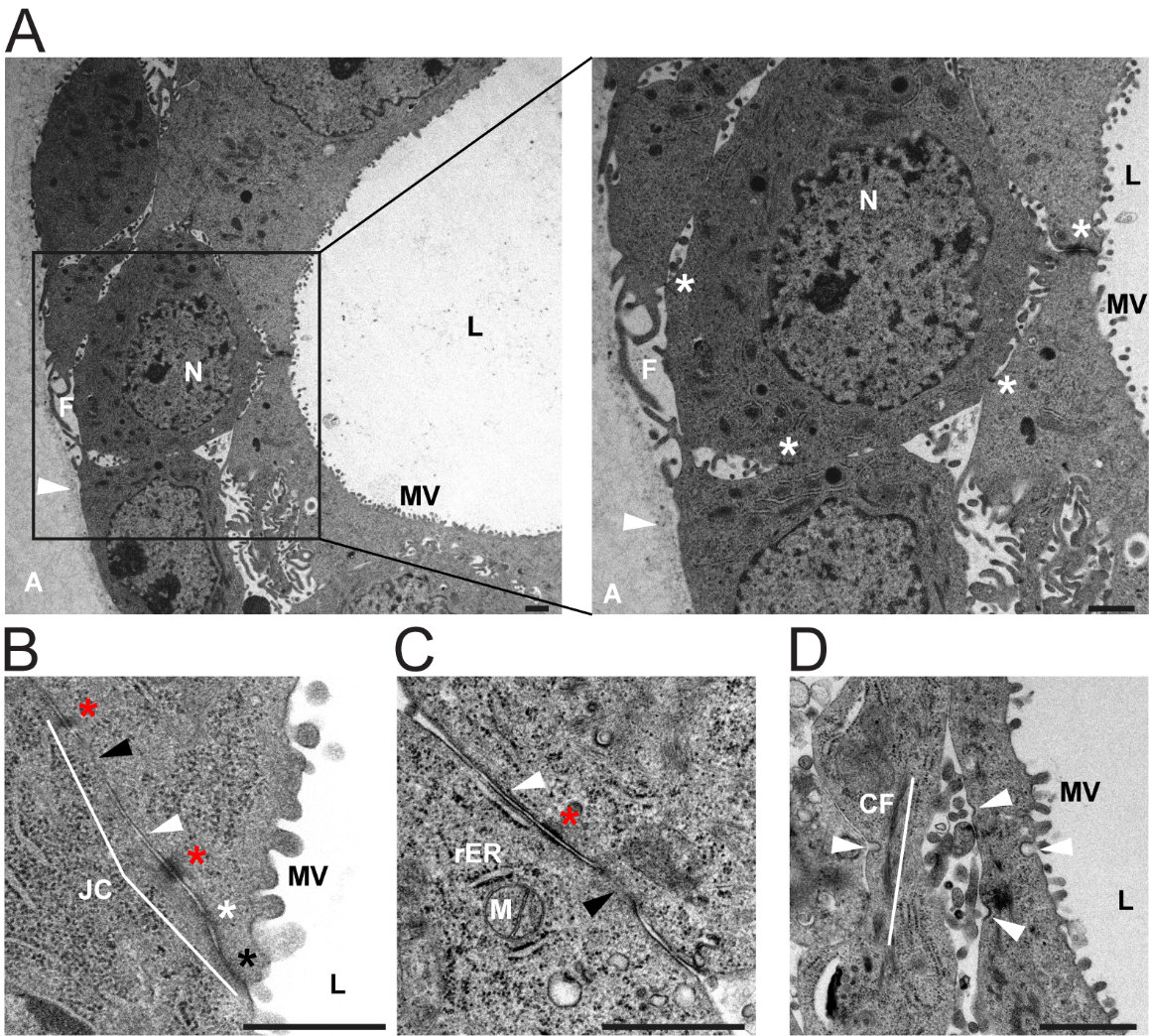

**S4 Figure. Kidney organoids display tubular epithelial polarity and complexity.** (A) Transmission electron microscopy of a cystic organoid with visible microvilli (MV) at the luminal side (L), a basal lamina without MV at the opposite basal side (arrowhead), filopodia (F), cell-cell contacts (asterisks) and surrounding agarose (A). (B) Detailed view showing an epithelial junctional complex (JC) between lateral membranes of neighboring cells. Shown are a tight junction (zonula occludens, black asterisk), an adherens junction (belt desmosome, zonula adhaerens, white asterisk), spot desmosomes (macula adhaerens, red asterisks) and closely aligned or nearly fused lateral membranes (white and black arrowhead, respectively). (C) Detailed view of a cell at the basal side. Shown are a spot desmosome (red asterisk) and closely aligned or nearly fused lateral membranes (white and black arrowhead, respectively). (D) Detailed view of endocytic events at apical and lateral membranes with clathrin-coated pits (diameter of around 100 nm, arrowheads). Data information: scale bars, 1 µm. Abbreviations; N, nucleus; M, mitochondrion; rER, rough endoplasmatic reticulum (dark ordered dots); CF, cytoskeleton filaments (actin, dark structures along the white line). In A, insets are enlarged on the right. Three independent replicates were examined.

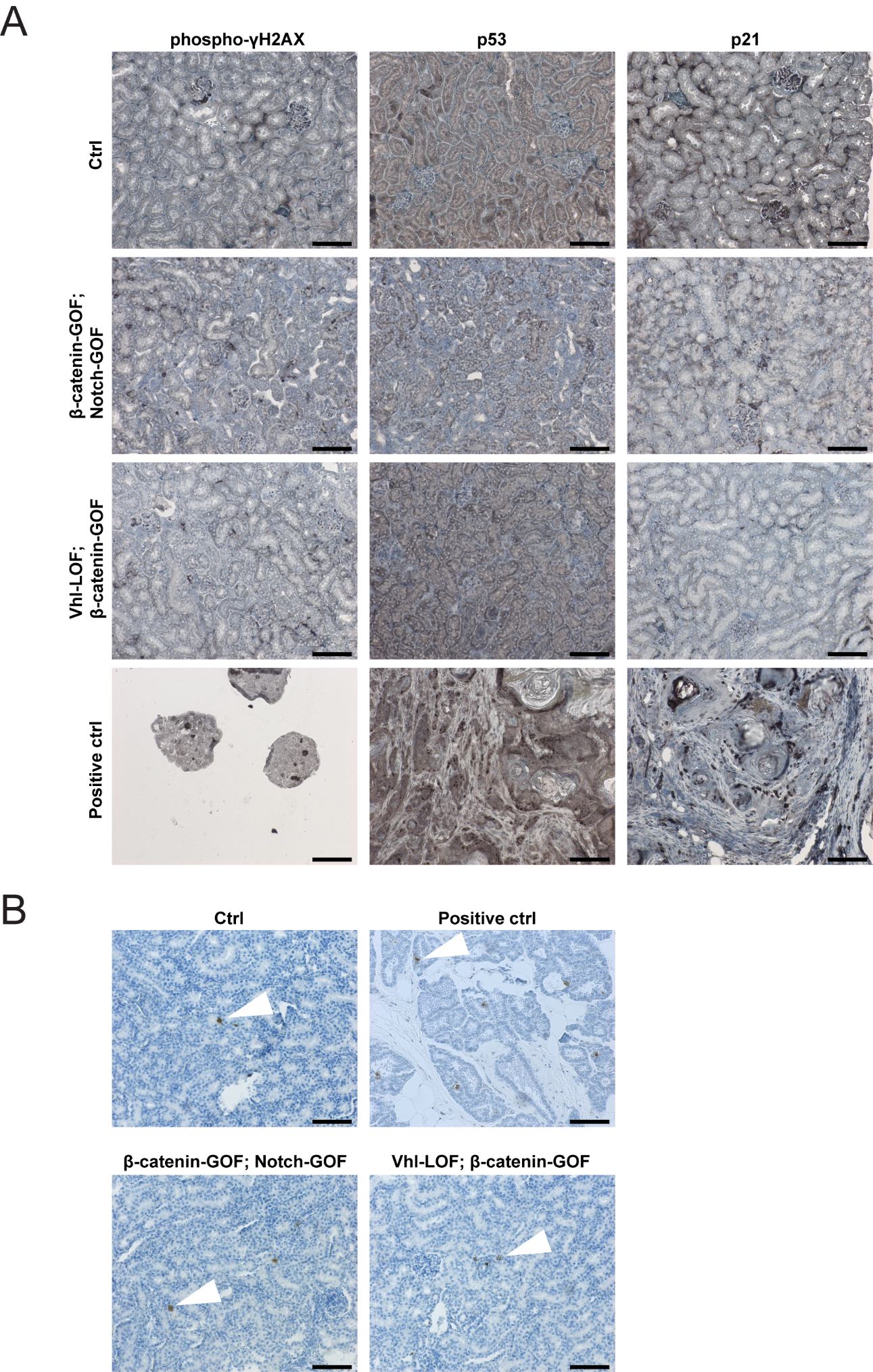

**S5 FIgure. No DNA damage, growth arrest or senescence and increased apoptosis was observed in mutant kidneys.** (A) Immunohistochemistry for nuclear phospho-γH2AX (left panel), p53 (middle panel) and p21 (right panel) in β-catenin-GOF; Notch-GOF and Vhl-LOF; β-catenin-GOF mutant cortical kidneys versus controls. (B) Immunohistochemistry for blunt ends of double-stranded DNA breaks in apoptotic cells (TUNEL assay) in β-catenin-GOF; Notch-GOF and Vhl-LOF; β-catenin-GOF mutant cortical kidneys versus controls. Data information: scale bars in A, 100 μm. Positive controls used; in the left panel, non-adherent spheres derived from a human colorectal carcinoma cell line LS174T with doxycycline-induced shRNA-mediated knockdown of *MLL1*; in the middle and right panel, mouse Pi3k-GOF; β-catenin-GOF; p53-GOF mammary gland tumors. Nuclei are counterstained with haematoxylin. Three independent replicates per line were examined. Scale bars in B, 100 μm. A positive control used (included in the kit), normal rat mammary gland 3-5 days after weaning of pups. Nuclei positive for blunt ends are marked by arrowheads. Nuclei are counterstained with haematoxylin. Three independent replicates per line were examined.

**S1 Video. Markers of tubular epithelial cells are expressed in 3D reconstructed kidney organoids.**
